# Synergistic Recruitment of UbcH7~Ub and Phosphorylated Ubl Domain Triggers Parkin Activation

**DOI:** 10.1101/344952

**Authors:** Tara E.C. Condos, Karen M. Dunkerley, E. Aisha Freeman, Kathryn R. Barber, Jacob D. Aguirre, Viduth K. Chaugule, Yiming Xiao, Lars Konermann, Helen Walden, Gary S. Shaw

## Abstract

The mechanism of activation and ubiquitin conjugation by the E3 ligase parkin is pivotal to understand the molecular pathology of early-onset Parkinson’s disease. Parkin is normally autoinhibited but is activated by the kinase PINK1 that phosphorylates parkin’s N-terminal ubiquitin-like (pUbl) domain and ubiquitin. How these alter the structure of parkin to allow recruitment of an E2~Ub conjugate to enhance ubiquitination is an unresolved question. We present the structure of an incoming E2~Ub conjugate with the phospho-ubiquitin bound C-terminus of parkin (R0RBR). We show the UbcH7~Ub conjugate is recruited by R0RBR parkin in the open state whereby conjugated ubiquitin binds to the RING1/IBR interface. Further, NMR experiments indicate there is re-modelling near the RING0/RING2 interface remote from the E2-binding site. This, and parkin phosphorylation lead to rapid reactivity of the RING2(Rcat) catalytic cysteine in parkin, needed for ubiquitin transfer. Parkin phosphorylation also leads to relocation and weak interaction of the pUbl domain with the RING0 domain that is enhanced upon E2~Ub recruitment indicating these events act synergistically to drive parkin activity.

---

Parkinson’s disease is the second most common neurodegenerative disease estimated to affect 1% of the population over 60 years of age (Tysnes & Storstein, 2017). The disease is believed to be a result of genetic predisposition or environmental factors (Corrigan *et al*, 1998) that lead to oxidative damage of mitochondrial proteins (Alam *et al*, 1997) and subsequent mitochondrial dysfunction (Schapira *et al*, 1990) and is characterised in patients by the loss of dopaminergic neurons in the substantia nigra of the midbrain (Hornykiewicz, 1966; Riederer & Wuketich, 1976). In addition to sporadic Parkinson’s disease there are also genetic forms of the disease that account for approximately 10% of all cases. In particular, mutations in the genes for *PARK2* and *PARK6* give rise to early-onset or autosomal recessive juvenile Parkinsonism (ARJP) forms of the disease that have similar symptoms including rigidity, bradykinesia and postural instability (Jankovic, 2008) but affect individuals at a much younger age. *PARK2* encodes the E3 ubiquitin-ligase parkin (Kitada *et al*, 1998) where mutations account for 50% of all ARJP cases. Along with the PTEN-induced kinase (PINK1) translated from *PARK6* these proteins use the ubiquitin degradation pathway to turnover damaged mitochondria and maintain mitochondrial homeostasis, especially under conditions of oxidative stress.

Parkin is a member of the RBR E3 ligase family that also includes the human homolog of Ariadne (HHARI) and HOIL-1 interacting protein (HOIP). These enzymes have a characteristic RBR motif comprising *R*ING1, in-*B*etween-RING and *R*ING2(*R*cat) domains that distinguish them from HECT and RING classes of E3 enzymes in terms of structure, mechanism and functionality. In particular, RBR E3 ligases incorporate a hybrid ubiquitination mechanism (Wenzel *et al*, 2011) whereby an E2 conjugating enzyme is recruited to the RING1 domain (similar to RING E3 ligases) and ubiquitin (Ub) is transferred from the E2~Ub conjugate to a catalytic cysteine in the RING2(Rcat) domain (similar to HECT E3 mechanisms) prior to labeling of a substrate lysine. RBR E3 ligases and RING E3 ligases have RING domains that are structurally similar and are expected to recruit E2 enzymes in a similar fashion (Budhidarmo *et al*, 2012), as recently shown in crystal structures of the RBR E3 ligases HHARI with UbcH7-Ub (Dove *et al*, 2017; Yuan *et al*, 2017) and HOIP with UbcH5b-Ub (Lechtenberg *et al*, 2016). However, a distinguishing feature of the HHARI and HOIP RBR E3 ligases is their ability to recognize an extended (“open”) form of the E2~Ub conjugate similar to that used by HECT E3 enzymes. This E2~Ub arrangement promotes a conformation susceptible to the transthiolation reaction needed to transfer the Ub cargo from the E2 enzyme to the catalytic cysteine of the RING2(Rcat) domain. Parkin on the other hand has been shown to function with a variety of E2 enzymes including UbcH7 and UbcH5b (Chaugule *et al*, 2011; Wenzel *et al*, 2011) although it appears that UbcH7 is the optimal E2 enzyme owing to its preference for ubiquitin transfer to cysteine, a requirement for RBR E3 ligases. While the conformation of the UbcH7~Ub conjugate during recruitment by parkin is unknown, it has been established that a cryptic Ub binding site within the RING1-IBR interface is only uncovered upon pUb binding to parkin and this has been proposed to help co-ordinate E2~Ub recruitment (Kumar *et al*, 2017).

All RBR E3 ligases identified to date, including parkin, appear to be uniquely regulated (Spratt *et al*, 2014; Walden & Rittinger, 2018). Parkin is normally autoinhibited by an accessory ubiquitin-like (Ubl) domain (Chaugule *et al*, 2011) that blocks both the E2 and cryptic ubiquitin sites. In addition, structures show that another accessory module, the RING0 domain, partially obscures the catalytic cysteine in the RING2(Rcat) domain protecting this site from Ub transfer. At least two steps have been identified for the activation of parkin both as a result of phosphorylation by PINK1. Under oxidative stress conditions PINK1 is activated and phosphorylates ubiquitin (pUb) near the outer mitochondrial membrane. This in turn helps recruit parkin to the membrane through binding of pUb to the RING1-IBR region of the E3 ligase (Kumar *et al*, 2017; Wauer *et al*, 2015; Sauve *et al*, 2015) and subsequent phosphorylation of parkin’s Ubl domain (Ordureau *et al*, 2014). These two events greatly stimulate ubiquitination activity (Kondapalli *et al*, 2012; Shiba-Fukushima *et al*, 2012; Kane *et al*, 2014; Kazlauskaite *et al*, 2014; Koyano *et al*, 2014) through an allosteric displacement mechanism of the pUbl domain from parkin (Kumar *et al*, 2015; Sauve *et al*, 2015). What is less clear is how parkin positions the E2~Ub conjugate enabling transfer of the Ub molecule to the RING2(Rcat) domain as a necessary step for catalysis. Current crystal structures of parkin show the proposed E2 binding site on the RING1 domain is greater than 50 Å from the catalytic site (C431) suggesting a significant conformational re-arrangement might be needed (Trempe *et al*, 2013; Riley *et al*, 2013; Wauer & Komander, 2013; Kumar *et al*, 2015; Sauve *et al*, 2015; Kumar *et al*, 2017; Wauer *et al*, 2015). A similar dilemma arises from recent structures of HHARI in complex with UbcH7-Ub that show the ubiquitin molecule is 47-53 Å from the its catalytic site (Dove *et al*, 2017; Yuan *et al*, 2017). Alternatively, structures of a HOIP:E2-Ub complex and a parkin:pUb complex raise the possibility of co-operation between multiple E3 ligase molecules to promote ubiquitin transfer (Lechtenberg *et al*, 2016; Kumar *et al*, 2017).

In this work we identify how the pUbl domain and E2~Ub conjugate co-operate to regulate parkin activity. We use NMR spectroscopy to determine a structure of pUb-activated parkin in complex with its biological UbcH7~Ub conjugate. The structure shows that UbcH7~Ub binds in an altered “open” conformation with its thioester linkage poised for Ub transfer to the catalytic cysteine of the RING2(Rcat) domain. We show that E2~Ub recruitment to parkin results in two distinct regions of NMR chemical shift changes; one set that is consistent with the E2~Ub binding site while a second set corresponds to residues near the RING0/RING2 interface, indicating this region is re-modeled during E2~Ub recruitment. In fully activated phosphorylated parkin in complex with pUb we have determined this corresponds to an increased exposure of the catalytic cysteine (C431) in the RING2(Rcat) domain. Further, we use NMR spectroscopy and hydrogen-deuterium exchange (HDX) experiments to show that the pUbl domain undergoes transient interaction with the RING0 domain that is enhanced upon E2~Ub recruitment by parkin. Overall our work provides a dynamic picture of parkin activation whereby PINK1 phosphorylation of parkin and E2~Ub recruitment co-operate to drive parkin activity.

## RESULTS

### The Parkin / UbcH7-Ub complex reveals an E2-Ub conformation poised for Ub transfer

Multiple high-resolution structures of parkin have provided significant biological insights into its E3 ligase function. Initial structures showed the compact nature of the C-terminal RING0-RING1-IBR-RING2(Rcat) domains (termed “R0RBR”) where the catalytic cysteine (C431) in the RING2(Rcat) domain is partially occluded (Trempe *et al*, 2013; Riley *et al*, 2013; Wauer & Komander, 2013). Structures of near full-length parkin have shown how the Ubl domain exerts its previously identified autoinhibitory effect (Kumar 2015, Sauve 2015) and details of partial E3 ligase activation have been shown in complexes with pUb (Kumar *et al*, 2015, 2017; Wauer *et al*, 2015). The association of pUb allosterically removes the pUbl domain from its interaction with the RING1 domain. Together these two phosphorylation events result in an increased affinity for UbcH7~Ub conjugate (Kumar *et al*, 2015) used by parkin to efficiently transfer ubiquitin to the RING2(Rcat) domain and propagate the ubiquitination reaction.

As a first step to identify how UbcH7~Ub transfers its Ub cargo to the catalytic cysteine within the RING2(Rcat) domain, we used NMR spectroscopy to examine how the E2~Ub complex is recruited to the R0RBR C-terminal region of parkin. These experiments utilized triple labeled (^2^H, ^15^N, ^13^C) human R0RBR parkin complexed with invisible ^2^H-labelled pUb and purified to homogeneity by size exclusion chromatography (R0RBR:pUb). NMR-based chemical shift perturbation experiments were monitored by ^1^H-^15^N TROSY spectroscopy between the selectively-labelled R0RBR:pUb complex and unlabelled UbcH7-Ub (isopeptide linked UbcH7 with Ub). These data show that many resonances in parkin R0RBR shift in the presence E2-Ub and undergo line broadening indicative of a nearly 70 kDa complex being formed (Supp. Fig. 1). Surprisingly, spectral changes were obvious in two distinct regions of R0RBR — one region where residues are mostly surface exposed and a second region where residues are largely buried (Figure 1A). The first area consists of resonances belonging to surface residues within the RING1, IBR and tether region indicative of the canonical E2/RING E3 binding interface. This binding region was confirmed through separate NMR experiments that monitored the individual binding of either unlabelled UbcH7 or Ub with selectively-labelled R0RBR:pUb. Reciprocal experiments using ^2^H-labelled R0RBR:pUb with ^2^H,^15^N-labelled UbcH7-Ub were monitored by chemical shift analysis of ^1^H-^15^N TROSY NMR spectroscopy (Supp. Fig. 2). In the absence of R0RBR, UbcH7-Ub is noted to be predominantly in the closed state based on chemical shift changes in helix H2 (K100-N113) of UbcH7 and G47-L50 in Ub, consistent with observations noted previously (Dove *et al*, 2016). Formation of a complex with R0RBR is indicated by line broadening and slow exchange binding kinetics for both UbcH7 and Ub moieties in the E2-Ub conjugate indicative of tight contact between both proteins with R0RBR:pUb. Notably we observe chemical shift changes in the UbcH7-Ub conjugate for UbcH7 (F22, V40, N43, N56 and Q106) and Ub (G47, Q49) that are hallmarks of UbcH7-Ub reverting to the open conformation upon binding to R0RBR. (Supp. Fig. 3). Together these observations show that the open state of UbcH7-Ub is induced through its recruitment by parkin.

**Figure 1.**
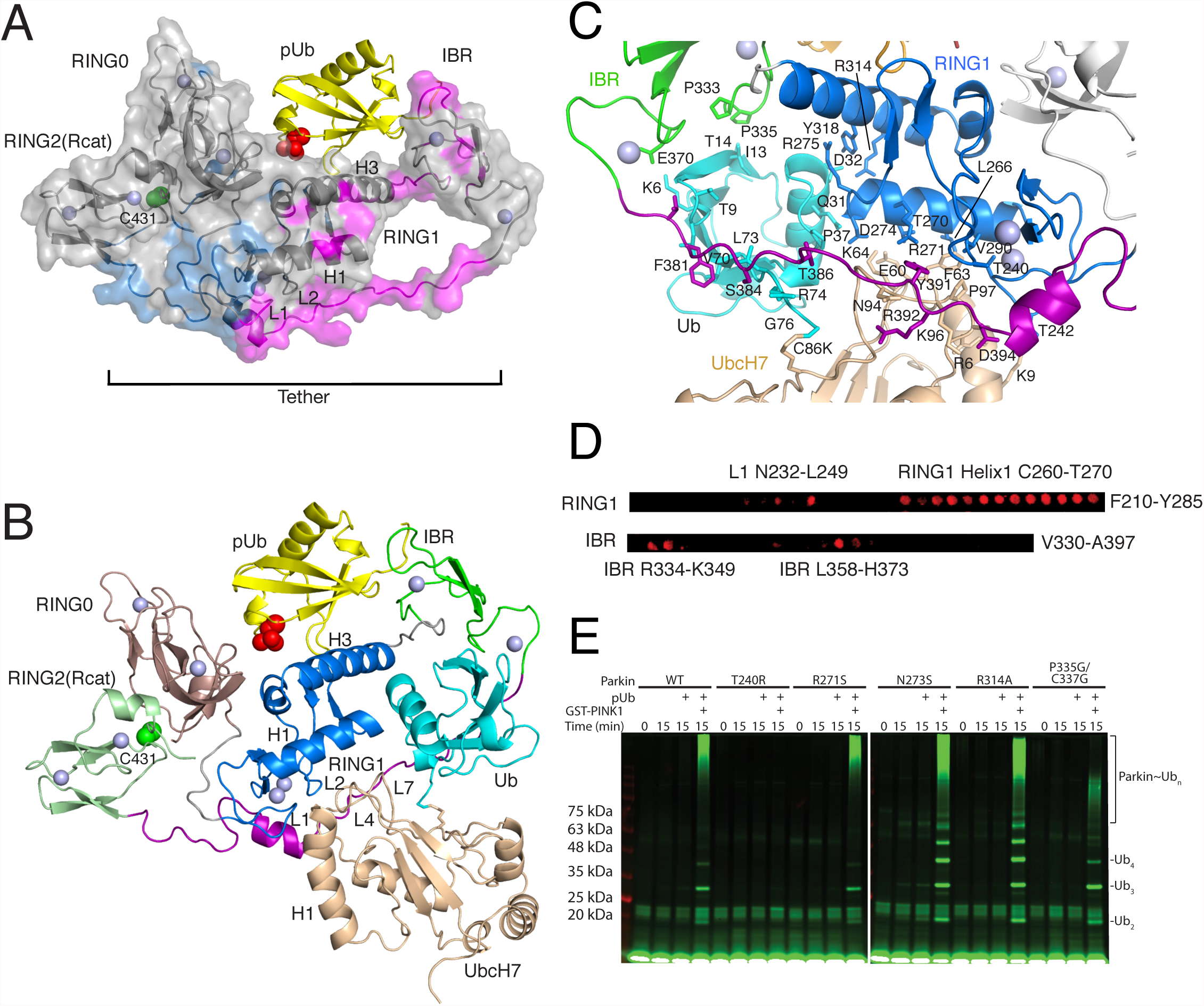
Model of the E2-Ub Conjugate bound to pUb-activated parkin. A. Two distinct surfaces are revealed on parkin upon binding of the UbcH7-Ub conjugate. Chemical shift perturbations were measured from ^1^H-^15^N TROSY spectra of ^2^H,^13^C,^15^N-labelled R0RBR parkin bound to unlabelled pUb in the absence and presence of one equivalent unlabelled UbcH7-Ub. Significant perturbations are modelled onto the surface of R0RBR bound to pUb (PDB 5N2W). The UbcH7-Ub binding site comprises the RING1 and IBR regions (magenta). An adjacent site composed of many buried residues was also observed (cyan). In this structure and accompanying data several sections of the tether region and some linkers were modelled in using Modeller so that chemical shift changes for residues not observed in crystal structures could be mapped. B. NMR-derived docked model of R0RBR:pUb with UbcH7-Ub. The lowest energy structure is shown although all 100 structures were very similar (RMSD 0.71 ± 0.1Å). The structure shows that UbcH7-Ub binds in an open conformation. UbcH7 uses canonical E2-RING E3 interactions that include residues in two loops in the RING1 domain (L1, L2) and two loops in the E2 enzyme (L4, L7) to stabilize the interaction. Ub binds to a RING1/IBR pocket. In this arrangement the distance between the G76 carboxylate and the catalytic C431 in the RING2(Rcat) is approximately 51 Å. C. Details of the UbcH7-Ub interaction with R0RBR:pUb. The isopeptide linkage between UbcH7 (C86K) and ubiquitin (G76) sits on the same side as helix 2 from the E2 enzyme. Specific residues that are at the interface for E2-Ub interaction with R0RBR parkin are highlighted. D. Peptide array for the RING1 and IBR domains of parkin showing selected residues are important for UbcH7-Ub recruitment. The arrays shown consist of RING1 (F210-Y285) and IBR (V330-A397) regions that are scanned using a series of 18-residue peptides that are shifted by two residues for each spot. The array was probed using Alexa-labelled UbcH7-Ub and imaged for fluorescence. E. Auto-ubiquitination assay for parkin using several substitutions observed near the interface with UbcH7-Ub. Assays were done in the absence and presence of pUb. Experiments with PINK1 were done by treating parkin:pUb with PINK1 for 30 minutes prior to adding other reagents needed for ubiquitination.

We used the chemical shift perturbation results to determine a model of UbcH7-Ub bound to R0RBR:pUb using HADDOCK (Figure 1B). In the model the location and orientation of the UbcH7 and Ub proteins with respect to R0RBR:pUb is similar in all 100 water-refined complexes and the best 20 complexes have a backbone RMSD of 0.71 ± 0.10 for R0RBR:pUb:UbcH7-Ub. As expected based on our NMR analysis, UbcH7-Ub takes on an open conformation in the complex where the UbcH7 moiety interacts mostly with the RING1 domain and the tether while Ub interacts with the RING1/IBR pocket created by pUb binding. The arrangement of the E2-Ub bound to R0RBR parkin more closely resembles the interaction of UbcH5b-Ub with HOIP than either of the structures for UbcH7-Ub with HHARI (Supp. Fig.4). Specifically, helix H1 (R6, K9) and loop L4 (E60, F63-K64) in UbcH7 contact the R0RBR loop L1 (T240, T242) and helix H1 (L266, T270, R271, D274) respectively in RING1 (Figure 1C). Unlike other E2/RING E3 ligase interactions the L7 loop (N94, K96, P97, A98) in UbcH7 sits between loop L2 (V290-G292) in RING1 and Y391, R392, V293 and D394 in the tether region. We noted that some of the largest chemical shift changes are in the tether region (Y391, R392, D394), yet multiple crystal structures of parkin in the presence and absence of its Ubl domain or bound to pUb alone show little structural change in this region. All structures to date show the tether blocks the UbcH7 binding site on the RING1 domain. However the current work (Supp. Fig. 5) and previous experiments (Spratt *et al*, 2013; Kumar *et al*, 2015) repeatedly show the regions before and after the REP helix in the tether are extremely flexible and sample a range of conformations. We interpreted these chemical shift changes to result from both direct E2 binding and an altering of the tether position to accommodate the E2 enzyme. Ub binding is governed predominantly by contacts from β1-L1-β2 (K6, L8, I13-T14), H1 (Q31, D32) and the following loop (P37), and its C-terminus (V70, L73, R74) to an R0RBR surface including β1 (P333, P335) and the C-terminus of the IBR domain (E370) and adjacent tether (V380, F381), RING1 helix H1 (D274) and the straightened RING1 helix H3 (R314, Y318) and in the tether (S384, T386). The Ub binding site in the IBR domain from UbcH7-Ub binding agrees well with potential ubiquitin-binding regions (UBR2, UBR3) inferred from crystallographic studies and supported through ubiquitination activity assays (Kumar *et al*, 2017). Further, the RING1 helix H1 site for UbcH7 and IBR site Ub were confirmed using a peptide array where we scanned 18-residue segments of the parkin RING1 and IBR sequences and probed with both fluorescently-tagged UbcH7 and UbcH7-Ub (Figure 1D). The orientation of the UbcH7-Ub in the structure poises the C-terminus of Ub for facile transthiolation by exposing the G76 carboxyl (isopeptide linkage) group towards the tether side of parkin. This arrangement suggests that nucleophilic attack by the catalytic C431 in the RING2(Rcat) domain would come from this direction (Figure 1C). The structure also shows that helix H2 in UbcH7, previously used to interact with Ub and favour the closed E2-Ub conformation, is exposed on the same side as the C86K-G76 linkage. The structure of parkin on the other hand is incompatible with ubiquitin transfer as the distance between ubiquitin (G76) and the catalytic cysteine (C431) in the RING2(Rcat) domain is greater than 50 Å.

In order to further test the parkin recruitment site for UbcH7-Ub observed in the structure a series ubiquitination assays were performed with ARJP and non-ARJP parkin variants in order to assess effects on parkin activity (Figure 1E). All assays were done in the absence and presence of pUb or with PINK1 preincubated with parkin and pUb to enable phosphorylation of the Ubl domain. As expected significant increases in ubiquitination were observed in the presence of both phosphorylation steps. ARJP variant T240R in the RING1 helix H1 shows significant decreases in activity due to disruption of interactions with F63 and P97 in UbcH7. The P335G/C337G substitution expected to disrupt one of the Zn-binding sites in the IBR domain that interacts with T13 and I14 of ubiquitin also had diminished activity. The R271S that is near the UbcH7 L4 loop had a minor decrease consistent with modification of that interaction. The activity of two other substitutions, N273S and R314A are more difficult to interpret. Although these neighbor the conjugated ubiquitin in the structure with R0RBR parkin the increased activity is more likely a reflection of a weakened interaction with the Ubl facilitating its phosphorylation and stimulating ubiquitination as previously observed (Sauve *et al*, 2015).

### UbcH7-Ub binding leads to a remodeled RING0/RING2 interface in parkin

In addition to the UbcH7-Ub binding site on parkin, analysis of chemical shift perturbation experiments of ^2^H,^13^C,^15^N-labelled R0RBR:pUb upon addition of UbcH7-Ub reveals many changes localized at the interface between the C-terminus of the tether region (A398-T414), RING0 (S145, F146), RING1 (Q252, R256, H257) and RING2(Rcat) (T415, E426, K427, N428, D464) (Figure 1A, 2A). Most of these residues form a cluster anchored by W403 of the REP element in the tether linker that is essential for packing the tether against the RING0/RING1/RING2(Rcat) core. The W403 side chain is also positioned nearby the C-terminal V465 at the RING0/RING2 interface in crystal structures although we were unable to resolve V465 in our spectra. Nevertheless, a W403A parkin variant has been shown to increase parkin activity (Trempe *et al*, 2013) proposed to be a result of the REP linker releasing interaction with RING1 and allowing access of the E2 to bind RING1 (Riley *et al*, 2013). However, the W403A variant may also result in the disruption of the RING0/RING2 interface. To test this, we compared resonances from a ^15^N-labelled R0RBR^W403A^ HSQC spectrum with those from an ^15^N-labelled R0RBR spectrum (Supp Fig. 6). In the R0RBR^W403A^ data many signals from residues that neighbor W403 are either undetectable or undergo significant chemical shift changes similar to that observed for UbcH7-Ub binding to R0RBR:pUb. Although we can not discount this region as a direct binding interface with UbcH7, the buried nature of many of these residues suggests the RING0/RING2(Rcat) interface is more likely re-modeled during E2~Ub interaction.

**Figure 2.**
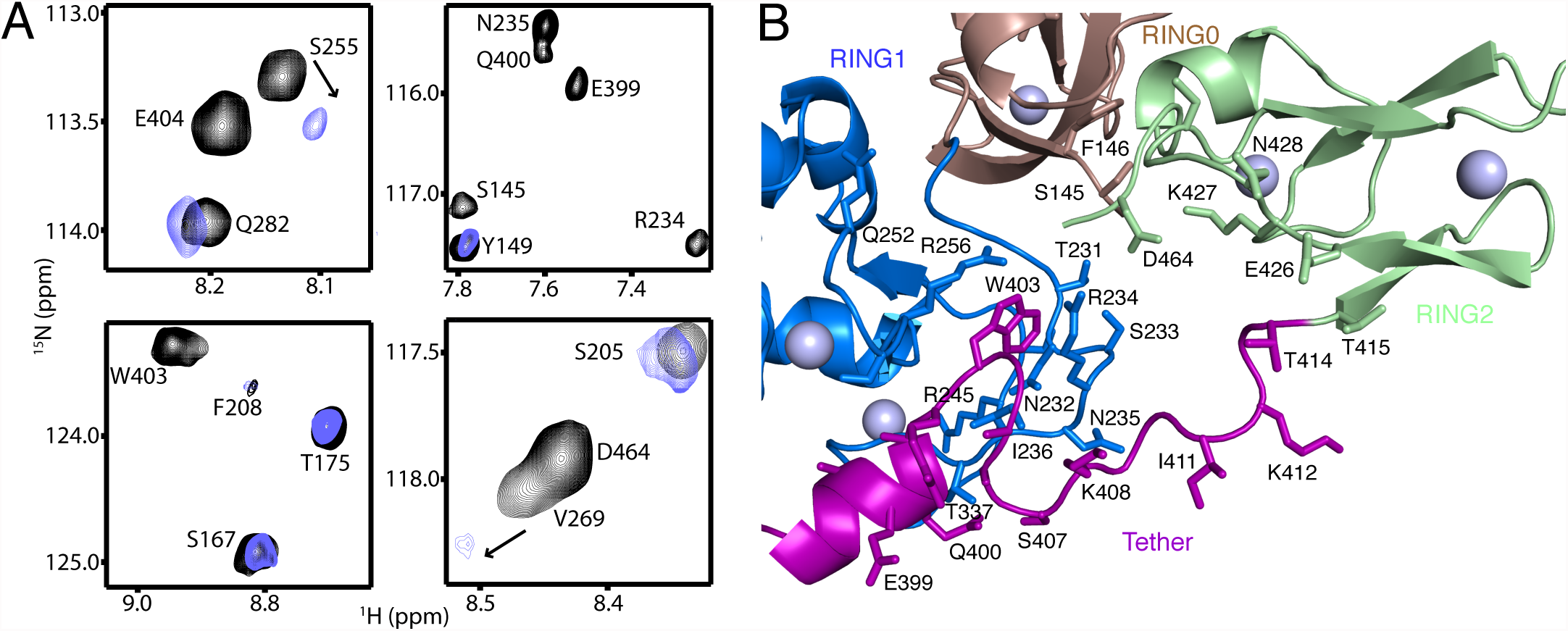
Remodelling of the RING0/RING2(Rcat) Interface induced by UbcH7-Ub Interaction. A. Selected regions from ^1^H-^15^N TROSY spectra of ^2^H,^13^C,^15^N-labelled R0RBR parkin bound to unlabelled pUb in the absence (black contours) and presence (blue contours) of one equivalent invisible ^2^H^,12^C^,14^N UbcH7-Ub. A large number of signals for residues including S145, N235, S255, E399, Q400, W403, E404 and D464 either shift or are absent in spectra with UbcH7-Ub. B. Details of the re-modelled region in parkin observed upon UbcH7-Ub binding. The figure shows selected residues in the RING0 (brown), RING1 (blue), tether (purple and RING2(Rcat) (pale green) interface that surround W403 and are most affected by UbcH7-Ub binding.

### UbcH7~Ub and pUbl work together to increase Cys431 reactivity

In its autoinhibited state, the Ubl domain blocks the E2 binding site in parkin resulting in minimal ubiquitination activity. Phosphorylation of parkin’s Ubl domain in combination with pUb binding lead to maximal ubiquitination activity of the E3 ligase through an allosteric displacement of the phosphorylated Ubl (pUbl) domain from its RING1-bound site. NMR and computational experiments show the phosphorylated Ubl (pUbl) domain samples a wide range of conformational space in this state (Caulfield *et al*, 2014; Aguirre *et al*, 2017). Our NMR experiments using R0RBR as a proxy for phosphorylated Ubl domain have allowed us to determine the UbcH7~Ub binding site within parkin as one requirement for ubiquitination. However, in the absence of the pUbl domain these experiments are deficient in establishing how the pUbl might work with an E2~Ub conjugate to achieve maximum activity. In particular all three-dimensional structures to date have been unable to show how the reactivity of the catalytic C431 in the RING2(Rcat) domain might be altered upon phosphorylation of the Ubl domain and upon presentation of the E2~Ub conjugate. In order to assess this, we examined the ability of parkin and R0RBR to form a non-hydrolyzable ubiquitin adduct using ubiquitin vinyl sulfone (UbVS). This approach tested the reactivity, and hence accessibility, of the catalytic C431 in the RING2(Rcat) domain during each step of the parkin activation cycle (Figure 3).

**Figure 3.**
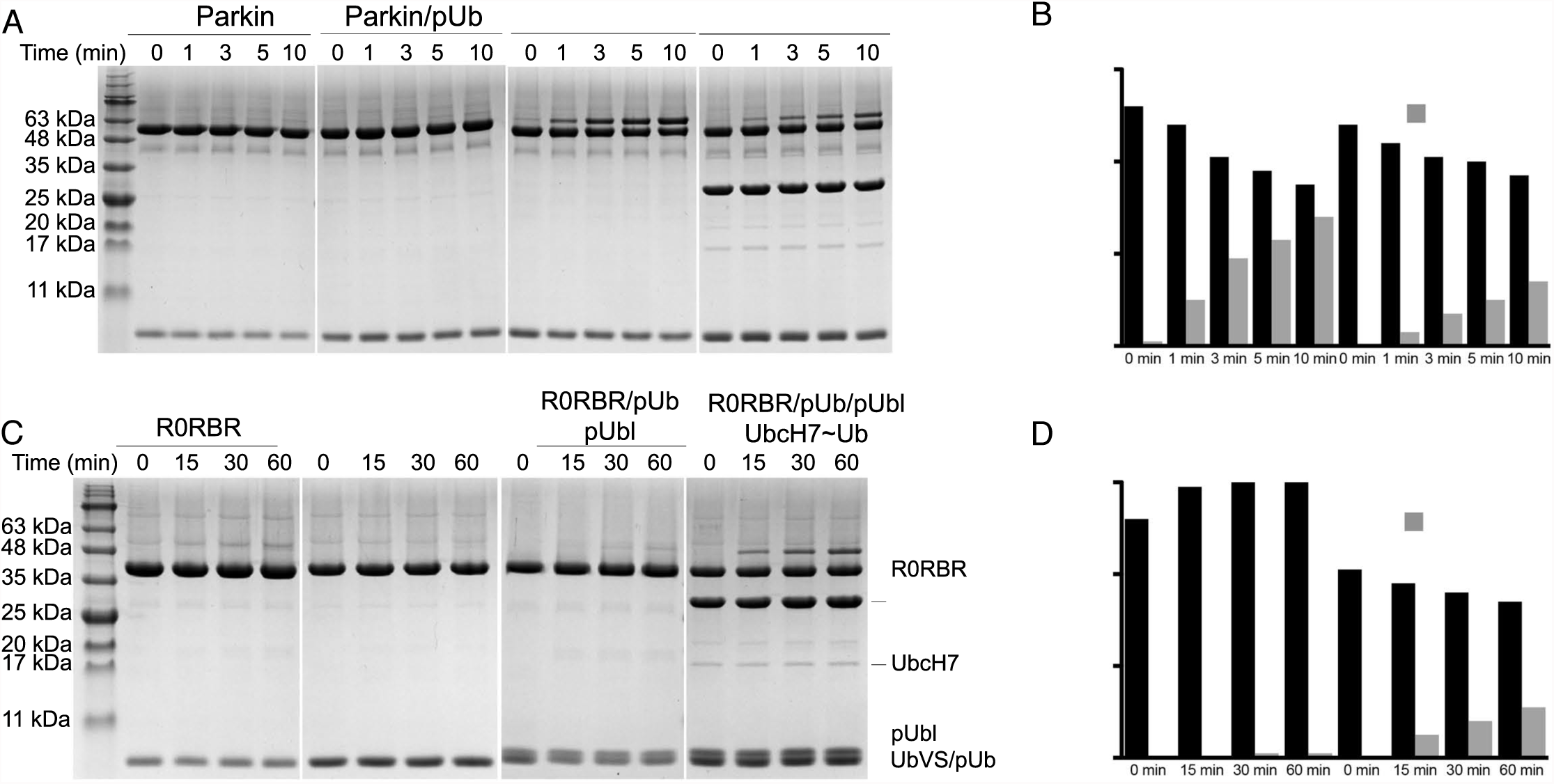
Ubl Phosphorylation and E2-Ub binding alter exposure of parkin’s catalytic cysteine. A. Exposure of the catalytic Cys431 is parkin as assessed by reaction with a UbVS probe. The different stages of activation using Parkin, Parkin:pUb, pParkin:pUb and pParkin:pUb in combination with an isopeptide linked UbcH7-Ub conjugate are indicated above each gel. Following addition of UbVS, samples were taken at the times indicated (0-10 minutes) and visualized by SDS-PAGE. B. Relative percentages of Parkin and the Parkin-Ub adduct as a function of time. Intensity percentages were calculated as a function of total intensity of Parkin-Ub, parkin and UbVS/pUb bands. C. Exposure of the catalytic Cys431 is R0RBR parkin as assessed by reaction with a UbVS probe. The different stages of activation using R0RBR, R0RBR:pUb, R0RBR:pUb:pUbl and R0RBR:pUb:pUbl in combination with an isopeptide linked UbcH7-Ub conjugate are indicated above each gel. Following addition of UbVS, samples were taken at the times indicated (0-60 minutes) and visualized by SDS-PAGE D. Relative percentages of R0RBR and the R0RBR-Ub adduct as a function of time. Intensity percentages were calculated as a function of total intensity of the R0RBR-Ub t, R0RBR and UbVS/pUb/pUbl bands.

As expected autoinhibited and pUb-bound parkin or R0RBR showed minimal reaction with UbVS (Figure 3) in agreement with previous three-dimensional structures and reactivity profiles that indicate the catalytic C431 is mostly occluded by neighbouring RING0 domain interactions. Remarkably, phosphorylated parkin activated by pUb shows rapid product formation with UbVS, visible even after 1 minute (Figure 3A, B). A similar reaction with R0RBR:pUb when the parkin pUbl domain is added *in trans* shows very little modification with UbVS even after 60 minutes (Figure 3C,D). The most logical explanation for these observations is that the pUbl domain is binding to another region in parkin that exhibits weaker affinity *in trans* than the local concentration afforded by pUbl in the intact protein. Interestingly, upon introduction of UbcH7-Ub, significant formation of the UbVS adduct was observed for both parkin and R0RBR. In the full-length protein we observe a reproducible lower conversion rate in the presence of the E2-Ub conjugate than in its absence (Figure 3B). This is surprising because in R0RBR the reactivity to UbVS is enhanced in the presence of UbcH7-Ub (Figure 3D). Given that the pUbl domain alone does not increase C431 accessibility *in trans*, this latter observation indicates that the E2-Ub conjugate likely works with the pUbl domain to re-model the RING0/RING2(Rcat) interface and increase reactivity of the catalytic C431 residue. Further, the decrease in C431 accessibility in the full-length protein may suggest that a further conformational change occurs due to E2~Ub binding that modifies availability of this site.

### Relocation and Re-binding of the pUbl Domain to the RING0 Domain

In order to test the potential relocation of the pUbl domain to a different binding site on parkin we examined ^1^H-^15^N HSQC spectra of the pUbl domain in the absence and presence of R0RBR:pUb and UbcH7-Ub. When invisible R0RBR:pUb is added to ^15^N pUbl we observe no significant chemical shift changes compared to ^15^N pUbl alone (Figure 4A) indicating that binding of pUbl to R0RBR is extremely weak or non-existent *in trans.* This observation agrees with UbVS experiments that showed pUbl is unable to enhance the reactivity of the catalytic C431 site in R0RBR (Figure 3C,D). Upon addition of UbcH7-Ub we observe small but measurable changes for the positions for several signals including pSer65, D62, Q63 and Q64 (Figure 4A). These chemical shift changes occur on the fast exchange timescale consistent with weak binding of the pUbl to R0RBR that is accentuated upon UbcH7-Ub recruitment by the E3 ligase. The potential re-location and re-binding of the pUbl to the C-terminus of parkin was also examined in full-length protein although the spectra are significantly more complicated due to the number of signals visible in ^1^H-^15^N HSQC spectra. In these data (Figure 4B) small changes in the positions of pSer65 and Q63 compared to the free, unbound pUbl domain occur consistent with a weak interaction of the pUbl domain in the absence of an E2-Ub conjugate. As with the R0RBR data the changes in the pUbl spectra are small and appear to occur on the fast chemical shift timescale consistent with weak binding. This indicates that the pUbl domain exists mostly in a dissociated state (Aguirre *et al*, 2017) but undergoes transient interaction with the remainder of the parkin protein.

**Figure 4.**
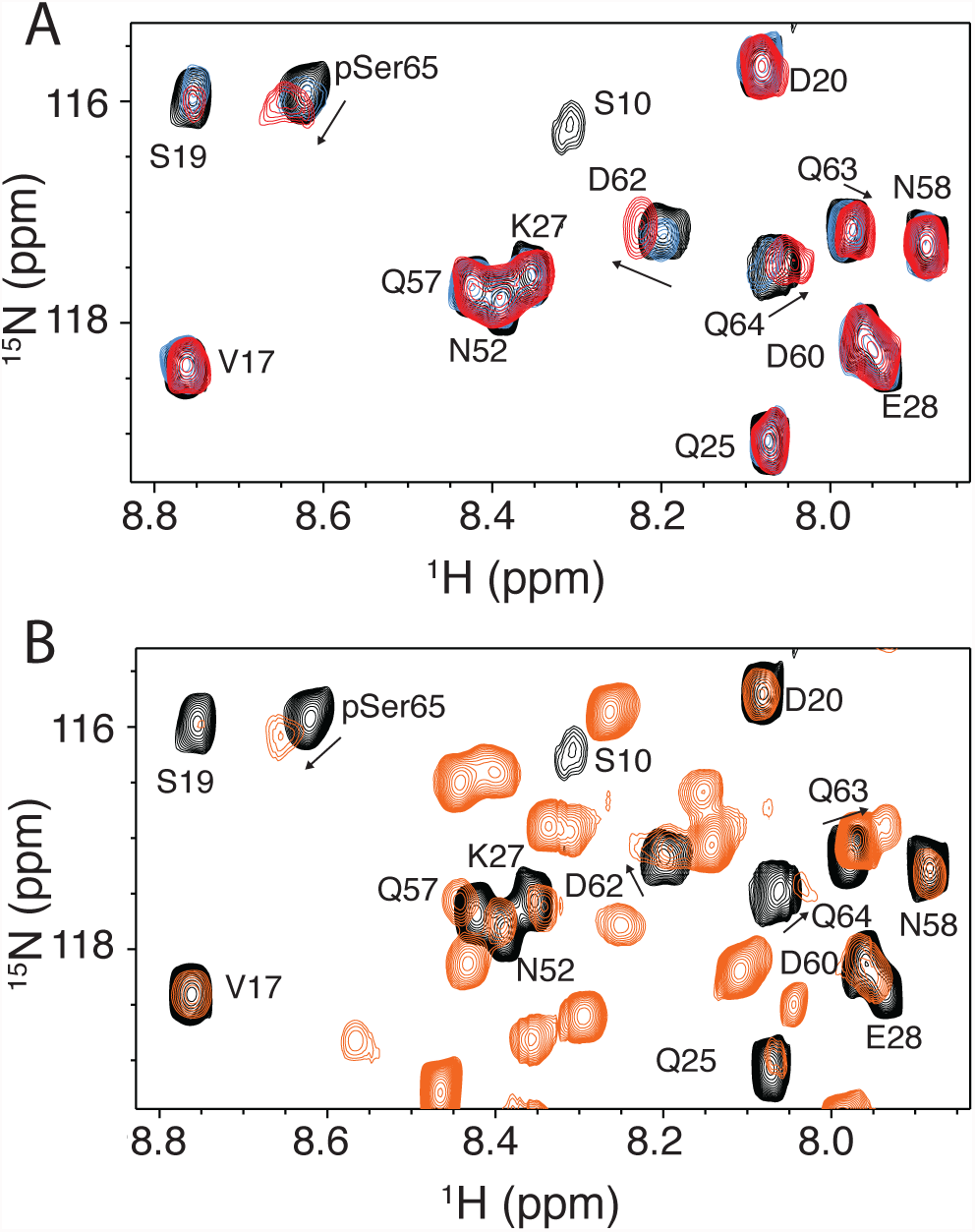
Weak interaction of Ubl with parkin after phosphorylation. A. ^1^H-^15^N HSQC spectra of ^15^N-labelled pUbl (black contours) and in the presence of an equimolar amount of unlabeled R0RBR:pUb (blue contours) shows little change. Upon addition of unlabelled isopeptide-linked UbcH7-Ub (red contours) small changes in chemical shift of pSer65, D62, Q63, Q64 are observed as indicated by arrows. B. ^1^H-^15^N HSQC spectra of ^15^N-labelled pUbl (black contours) compared to full-length, ^15^N-labelled pParkin: pUb (unlabelled) examined using a ^1^H-^15^N CPMG, T2-filtered HSQC (orange contours) (Aguirre *et al*, 2017). In the spectrum of full-length parkin many resonances arising from flexible loops not found in the pUbl domain are evident and are not labelled for clarity. Residues that experience small chemical shift changes are indicated by arrows.

To identify the possible location of interaction of the pUbl domain with parkin we used hydrogen-deuterium exchange experiments measured by mass spectrometry (Figure 5, Supp Fig 7). We compared the increase in deuteration of phosphorylated parkin in complex with pUb (pParkin:pUb) to autoinhibited parkin. In pParkin:pUb we observe a general increase in deuteration consistently across most of the pUbl domain indicating that upon dual phosphorylation this domain is more exposed compared to its autoinhibited state where it is protected through interaction with the RING1, IBR and tether regions of parkin. Coincident with this, increased exposure was noted at the IBR/tether junction (C365-A379) which has now been released from pUbl interaction. As expected, a series of peptides covering regions of the RING0 (147-155, F209-A225) and RING1 (F277-L283, I298-E309, Y312-E322, V324-L331) show slower exchange in the pParkin:pUb state due to protection of this region by pUb as revealed in multiple crystal structures.

**Figure 5.**
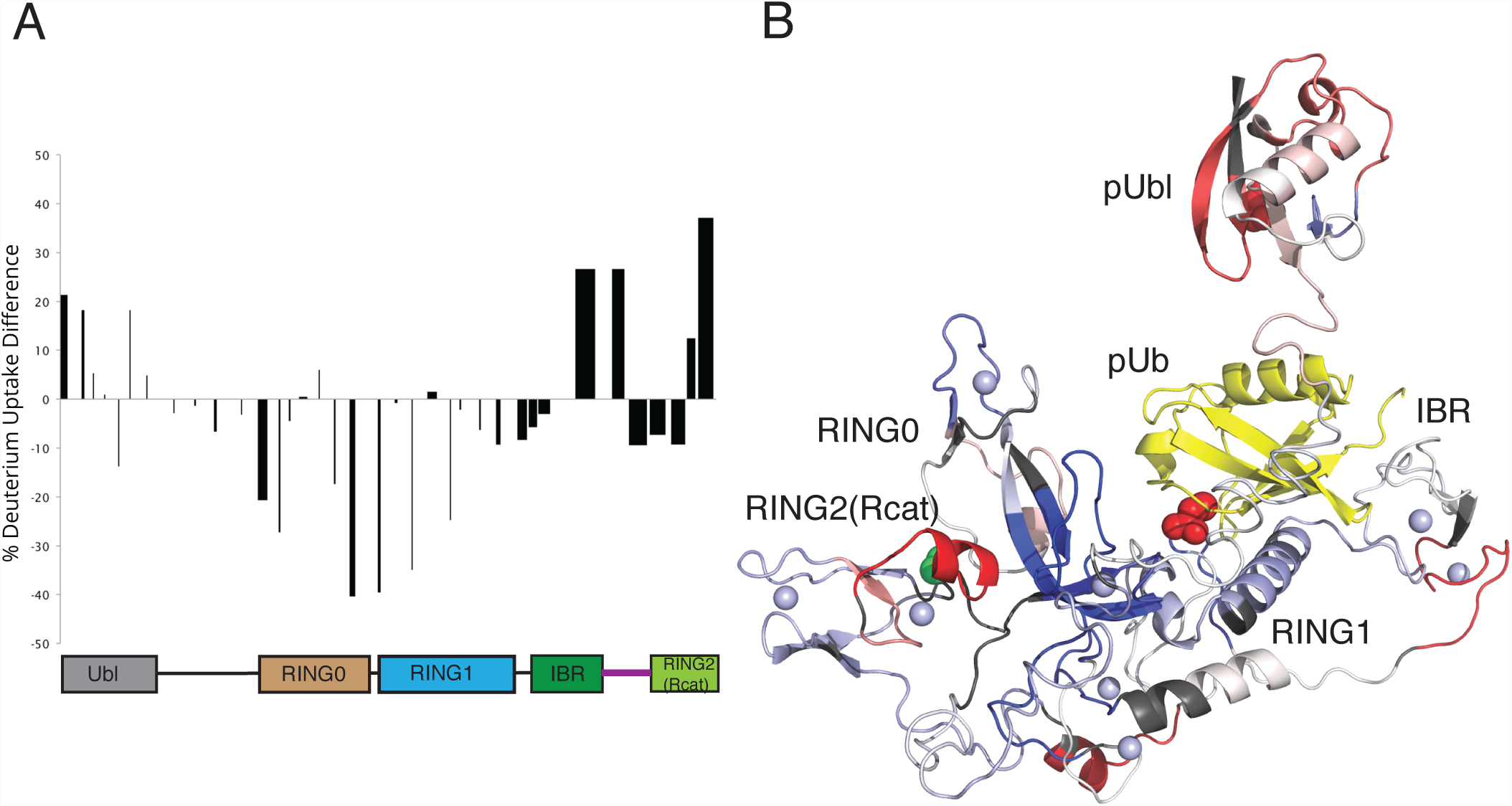
Identification of a pUbl binding site and exposure of RING2(Rcat) C-terminus. A. HDX plot showing the differences for deuterium uptake between auto-inhibited parkin and activated pParkin:pUb state at the 1 minute time point. The width of the bars shown represent the lengths of the peptides observed and measured while the heights represent the relative difference in hydrogen-deuterium exchange. The peptide length is shown through the width of the column on the graph. Positive bars indicate an increase of deuterium uptake in the pParkin:pUb state indicating greater exposure to solvent, while a negative change indicates a decrease in deuterium uptake indicating the site is more protected. The domain structure of parkin is shown below the data to indicate the location of peptides. B. Relative HDX differences are mapped to the structure of parkin:pUb (PDB 5N2W). Regions of the protein not visible in crystal structures were added to complete the structure. The location of the pUbl domain is shown in a dissociated state from the remainder of the protein in agreement with analytical sedimentation velocity experiments (Aguirre *et al*, 2017) and for clarity to map exchange sites.

Some of the largest differences in exchange between pParkin:pUb and autoinhibited parkin occur near the RING0/RING2(Rcat) interface. These include peptides with decreased deuteration uptake in the RING0 domain (Y147-Q155, R156-L162, F209-A225, H227-T237) that form a near contiguous surface on this domain (Figure 5B). This observation would be consistent with the pUbl domain re-locating and binding to this surface. This region also includes three basic residues (K161, R163 and K211) in the RING0 that show attenuated parkin autoubiquitination when substituted (Wauer *et al*, 2015). Two of these sites (K161N, K211R, K211N) are locations of ARJP substitutions. In addition, a minority of parkin crystal structures show a bound sulphate ion in this region. Altogether these data suggest a possible site on the RING0 domain for a weak pUbl interaction. Interestingly, the pParkin:pUb state also has two peptides with enhanced exchange at the C-terminus (E444-G450, E452-W462) of the RING2(Rcat) domain. All three-dimensional structures of parkin to date reveal that the E452-W461 region, which includes a short two-turn helix, packs against the RING0 domain β7 sheet consistent with slower amide exchange in the autoinhibited state. In pParkin:pUb exchange is more rapid signifying this region is more exposed. These two observations signify that interaction between the C-terminus of the RING2(Rcat) domain and the RING0 domain is more open, possibly accommodating pUbl interaction by the RING0 domain. This structural change would be expected to increase the reactivity of the catalytic cysteine C431 as observed in our UbVS experiments (Figure 3A,B).

## DISCUSSION

Although parkin was originally thought to be a constitutively active enzyme, it is now known to regulate its activity through intramolecular domain-domain interactions, and binding to effectors (Chaugule *et al*, 2011a; Kumar *et al*, 2015; Sauve *et al*, 2015). In particular, PINK1 regulates parkin activity through phosphorylation of the Ubl domain, and of ubiquitin itself, which then acts as an effector (Figure 6). Multiple structures of autoinhibited parkin reveal that the E2-binding site is blocked, however, static crystal structures have shown that binding to pUb does not render the proposed E2 binding site accessible. Similarly, pUb binding to parkin decreases the affinity of R0RBR for the Ubl domain, but is not sufficient to dislodge the Ubl domain in the crystal (Kumar *et al*, 2015, 2017). A major caveat of such studies is that the long linker between the Ubl domain and RING0 hinders crystallization and is removed, thus reducing flexibility. However, the linker plays a role in parkin function because different truncations have altered activities, and modelling analysis shows that the linker promotes dissociation of the Ubl domain (Caulfield *et al*, 2014; Kumar *et al*, 2015; Aguirre *et al*, 2017). Nevertheless, comparison of crystal structures in the absence/presence of the Ubl domain reveal reorganization of residues between the RING0/RING1 interface (Kumar *et al*, 2015) and flexibility of the IBR domain thought to optimize parkin for E2~Ub engagement (Figure 6A,B). Furthermore, dynamic analysis of full-length phosphorylated parkin bound to pUb shows increased flexibility of the Ubl domain (Aguirre *et al*, 2017) (Figure 6C). Our data show that in addition to pUb binding, both phosphorylation of the Ubl domain and binding of the loaded E2 lead to increased reactivity of parkin’s catalytic cysteine. Furthermore, mutation of W403, a hydrophobic residue that pins several domains together, has previously been shown to support parkin labelling with an E2-based activity probe, even in the absence of Ubl domain phosphorylation (Pao et al., 2016). We find that mutation of W403 gives rise to a similar remodeling as binding of E2-Ub, suggesting that E2-Ub binding induces a conformational change in the W403 cluster (Figure 6D), rather than W403 needing to remodel to allow E2-Ub engagement. The dynamics of the interaction between pParkin:pUb and E2~Ub show that the donor Ub is in the open conformation favoured by RBRs (Dove *et al*, 2016), and that it is bound in the cryptic Ub site exposed upon pUb binding (Kumar *et al*, 2017). Furthermore, we find that the catalytic domain of parkin has enhanced reactivity in the presence of both E2~Ub and pUbl, indicating synergy between these events. Interestingly, phosphorylation of the Ubl, binding of the E2~Ub, and dynamics near the RING0/RING2(Rcat) interface, lead to a weak secondary docking site for the pUbl on the RING0 domain (Figure 6C,D). This secondary interaction site likely includes a previously identified (Wauer & Komander, 2013) basic patch formed by K161, R163, K211 sites of two ARJP substitutions. Mutation of these residues renders parkin unreactive with an E2-based activity probe, consistent with a requirement for pUbl interaction (Pao et al., 2016). The fast exchange observed in current NMR experiments between the pUbl domain and RING0 domain suggests that this interaction is short-lived. Addition of the pUbl domain to R0RBR:pUb does not allow for the catalytic cysteine to react with a ubiquitin probe. This suggests that pUbl binding alone to R0RBR parkin is not sufficient to drive the conformational changes required, and that E2~Ub binding is necessary for full activation (Figure 6E). This agrees with several studies that have previously demonstrated a loss of pUbl binding to R0RBR:pUb (Kumar *et al*, 2015; Sauve *et al*, 2015; Wauer *et al*, 2015). Taken together, the evidence suggests that there is a rapid association/dissociation of the pUbl domain with R0RBR:pUb, and we speculate that this may be to ensure parkin activity only in the presence of the loaded E2~Ub. Our study suggests that parkin activity as stimulated by PINK1 is tightly controlled, and proceeds through several intermediate steps. Understanding those steps in detail will be essential to targeting this important enzyme for modulation during the pathogenesis of PD.

**Figure 6.**
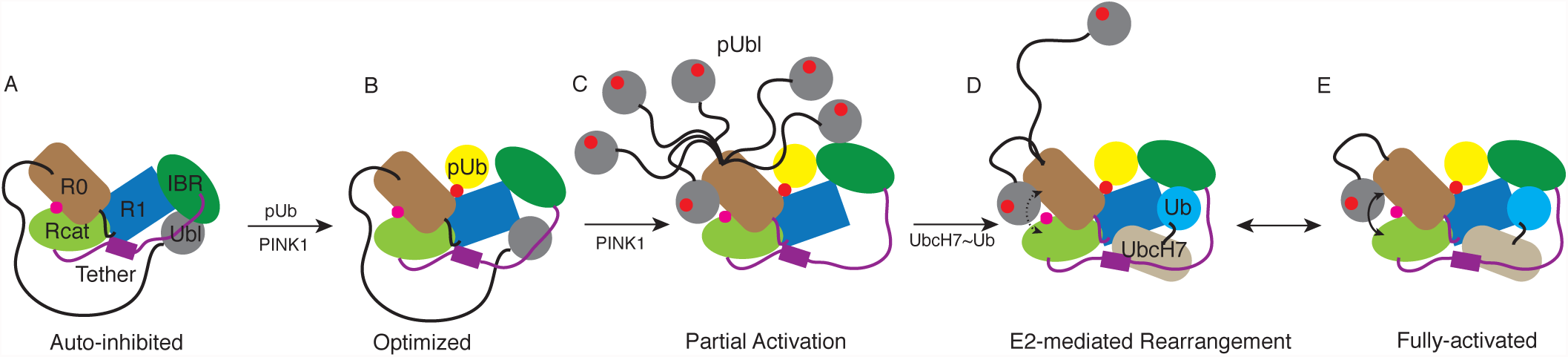
Model of parkin activation by combined pUbl and E2~Ub interactions. Highly controlled activation of parkin is regulated by multiple steps. (A) Autoinhibited state whereby the Ubl domain masks the E2 binding site on the RING1 (R1) domain as described by Chaugule *et al* (2011) and supported through crystallographic studies. (B) Optimization step controlled by PINK1 phosphorylation of ubiquitin (Ub) and subsequent binding of pUb to the RING1/IBR interface resulting reorganization of the RING0/RING1 interface and movement of the IBR domain. Steps (A) and (B) have been previously described. (C) PINK1 phosphorylation of the Ubl domain causes its dissociation from the RING1 binding site allowing it to sample a large conformational space in solution including weak binding to the RING0 domain favouring an uncovering of the catalytic cysteine C431. (D) E2~Ub binding to the RING1/IBR domains making use of a cryptic ubiquitin binding site uncovered through dissociation of the pUbl domain. In both (C) and (D) re-modelling of the RING0/RING1 interface with the tether region occur based on NMR chemical shift data and heteronuclear NOE experiments. (E) Fully activated state of parkin utilizes synergistic binding of pUbl domain to the RING0 domain and E2~Ub binding to maximize accessibility to the catalytic cysteine.

## METHODS

### Protein Constructs and Purification

Full-length human parkin (1-465), Ubl (1-76), R0RBR (141-465), *Drosophila melanogaster* RING2 (410-482) and other parkin variants were expressed and purified as described previously (Chaugule *et al*, 2011a; Spratt *et al*, 2013). Briefly, His-smt3-parkin constructs were expressed in BL21(DE3) cells at 37°C to an OD_600_ of 0.8. Expression was induced at 16°C with 25 µM IPTG for parkin, 0.1 mM IPTG for R0RBR, and 0.5 mM IPTG for Ubl or RING2 for 18 hours. All growths, except the Ubl domain, were supplemented with 0.5 mM ZnCl_2_. Purification utilized an initial HisTrap FF column followed by Ulp1-cleavage at 4°C, a second HisTrap FF column and final Superdex 75 10/300 size-exclusion chromatography. Selectively ^2^H,^13^C,^15^N-labelled R0RBR:pUb or R0RBR and selectively ^2^H^,13^C^,15^N or ^2^H,^15^N-labelled Ub, UbcH7 or UbcH7-Ub were expressed and purified as previously described (Kumar *et al*, 2015).

His-tagged Uba1 was expressed in BL21(DE3)CodonPlus-RIL cells at 37°C to an OD_600_ of 0.8. Expression was induced with 0.5 mM IPTG at 18°C for 12 hours. His-tagged Uba1 was purified on a HisTrap FF column by washing with 50 mM Tris, 200 mM NaCl, 250 μM TCEP, 25 mM imidazole, pH 8.0 buffer and then washing with 14% of elution buffer that contained 250 mM imidazole. The His-tagged Uba1 was then eluted with 100% elution buffer and stored in aliquots at -80°C for ubiquitination assays.

His-TEV-tagged human UbcH7^C17S,C86K,C137S^ was expressed in BL21(DE3)CodonPlus-RIL cells at 37°C to an OD_600_ of 0.8 and expression was induced with 1 mM IPTG at 30°C for 18 hours. UbcH7 was purified on a HisTrap FF column, cleaved at 4°C overnight and purified on a second HisTrap FF column. His-tagged Ub was expressed in BL21(DE3)CodonPlus-RIL cells.

Complexes that contained 1:1 R0RBR:pUb were formed using a 1.5 fold excess pUb compared to R0RBR in 20 mM Tris, 75 mM NaCl, 250 µM TCEP, pH 8 buffer. The 1:1 complex mixture was purified on a Superdex 75 10/300 size-exclusion column to ensure excess pUb was not present in samples for NMR studies.

### Protein Phosphorylation

Phosphorylation of Ub, Ubl and parkin were done using purified *Pediculus humanus* PINK1 (126-575) as described previously (Kumar *et al*, 2015; Aguirre *et al*, 2017). For pUb and pUbl (1-76) typically 10 µM PINK1 was sufficient to stoichiometrically phosphorylate either 100 µM Ub or Ubl in 1 hour at 24°C. For parkin, typically 75 µM PINK1 was sufficient to phosphorylate 150 µM parkin for 3.5 hours at room temperature. Reactions were visualised by Phos-tag gel. PINK1 was removed using a GSTrap FF column. Phosphorylated proteins were purified using a Superdex 75 10/300 size-exclusion column and confirmed by mass spectrometry.

### Synthesis of UbcH7-Ub isopeptide-linked conjugate

UbcH7-Ub isopeptide-linked conjugate was synthesized using an optimised version of the protocol of Plechanovová *et al* (2012). Briefly, 200 µM His-tagged Ub, 400 µM UbcH7^C17S/C86K/C137S^, 25 µM non-cleavable His-tagged Uba1 and 10 mM Mg^2+^/ATP were incubated together in 50 mM CHES, 150 mM NaCl, pH 9.0 buffer at 37°C for 6-16 hours to form approximately 60% UbcH7-Ub isopeptide-linked conjugate based on SDS PAGE analysis. The resulting mixture was passed through a HisTrap FF column to eliminate unconjugated UbcH7. The eluted His-tagged proteins were TEV-cleaved overnight at 4°C. The mixture was purified on a second HisTrap FF column to eliminate non-cleavable His-tagged Uba1. The remaining UbcH7-Ub was separated from unreacted Ub using a HiLoad Superdex 16/60 size-exclusion column.

### NMR experiments

All NMR data were collected at 25°C on a Varian Inova 600 MHz NMR spectrometer equipped with a triple resonance cryogenic probe and z-field gradients. Samples were prepared in 25 mM HEPES, 50 mM NaCl, 500 µM TCEP, pH 7.0 buffer with 10% D2O (v/v) using DSS as an internal reference and imidazole to monitor pH. ^1^H-^15^N HSQC were collected in TROSY mode (Pervushin *et al*, 1997) to follow amide backbone chemical shift perturbations. ^1^H-^13^C HMQC spectra (Tugarinov *et al*, 2004) were collected to monitor chemical shifts of Ub side chain methyl groups. ^1^H-^15^N TROSY spectra were collected using different combinations of ^2^H,^13^C,^15^N-labelled and ^2^H,^15^N-labelled R0RBR:pUb or R0RBR with ^2^H,^13^C,^15^N or ^2^H,^15^N-labelled pUbl, Ub, UbcH7 or UbcH7-Ub. Chemical shift perturbation measurements for amide backbone resonances were calculated using Δδ=((ΔδH)^2^+(ΔδN/5)^2^)^0.5^ and for side chain methyl groups using Δδ=((ΔδH)^2^+(ΔδC/3.3)^2^)^0.5^. All data was processed using 60°-shifted cosine bell-weighting functions using NMRPipe and NMRDraw (Delaglio *et al*, 1995), and was analysed using NMRViewJ (Johnson & Blevins, 1994).

### Ubiquitin~Vinyl Sulfone Reactions

Individually purified proteins were acquired as described above. The final concentration of each component was 10 µM in a final volume of 45 µL (R0RBR and Rcat) or 55 µL (parkin and 77C) in 50 mM HEPES, 50 mM NaCl, pH 8.0. Time started when the UbVS was added to the reaction at 37°C. 10 µL were removed at each time point and the reaction was quenched with 3xSDS sample buffer. 16.5% SDS-PAGE gels were run and stained with Coomassie Blue. Gels were imaged on a BioRad ChemiDoc XRS+. Band intensities of parkin~Ub, parkin, UbVS/pUb/pUbl were measured using ImageJ software (Schneider *et al*, 2012). The percent contribution per band was calculated from the combined intensity in each lane.

### Ubiquitination Assays

All reactions were performed at 37°C contained purified 1 µM WT or substituted parkin, 0.5 µM UbcH7, 0.1 µM Uba1, 4 µM Ub and 0.5 µM Ub^800^ in 5 mM MgATP, 50 mM HEPES (pH 7.5). pUb was added to 0.5 µM when needed. To induce *in situ* phosphorylation, 0.01 µM of purified GST-PINK1 was added to the parkin/pUb/ATP samples 30 min before initiating ubiquitination. The ubiquitination reactions were quenched with 3xSDS sample buffer and 1 M DTT. 4%-12% Bis-Tris gradient gels (Thermo Fisher Scientific) were used with MES running buffer (250 mM MES, 250 mM Tris, 0.5% SDS and 5 mM EDTA, pH 7.3). Fluorescence intensity at 700 and 800 nm was measured using an Odyssey Imaging system (LiCor).

### HDX Mass Spectrometry

Deuteration of proteins occurred at 20°C ± 1 °C in 90% D_2_O and 10% H_2_O with 50 mM HEPES, 100 mM NaCl and 250 μM TCEP at pH 7.0. Protein concentrations were 1 μM. UbcH7-Ub conjugate concentration was 0.5 μM within the pParkinp-pUb complex for a 1:2 ratio respectively. 100 μL aliquots were removed at time points between 15 seconds and 10 minutes following deuteration. Aliquots were quenched with ice-chilled 10% HCl in H_2_O to reach a pH of 2.3 and then flash frozen in liquid nitrogen. Zero time point (*m_0_*) controls were created by adding ice-chilled D_2_O to an ice-chilled protein sample under quench conditions (pH 2.3) and flash frozen. Fully exchanged controls (*m_100_*) were also created by exposing the proteins to D_2_O at pH 2.3 and heated to 70°C with a water bath for 8 hours. These samples were then flash frozen in liquid nitrogen. Aliquots were thawed to approximately 0° C and injected into a Waters H DX nanoAQUITY HPLC system. Online digestion of the proteins was performed with a POROS pepsin column at 15°C. Resulting peptides were trapped and analyzed on a Waters BEH C18 column at 0°C using a water/acetonitrile with 0.1% formic acid gradient at 40 μL/min. Peptide masses were measured using a Waters Synapt G2 Q-TOF mass spectrometer. Peptides were identified through MS/MS. Resulting peptides were analyzed with Waters Dynamix 3.0. Deuteration is expressed here as percent deuteration uptake where *mt* is the centroid mass at time *t* and *m_0_* and *m_100_* are described above according to the following equation.

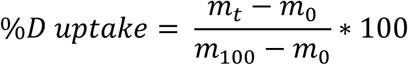

### Model Determinations for UbcH7-Ub binding to R0RBR:pUb

Interacting residues were identified from NMR experiments (described above) and defined as those amides that shifted greater than the average + one standard deviation and had greater than 20% side chain accessible surface area. Additional interacting residues were determined through mutagenesis studies that showed reduced binding and/or ubiquitination by parkin. Passive residues were defined according to the HADDOCK protocol as residues that neighbored active residues had greater than 20% side chain accessible surface area and had a noticeable chemical shift change.

The UbcH7-Ub conjugate was docked to R0RBR:pUb using HADDOCK (Dominguez *et al*, 2003) using NMR data as restraints. Starting coordinates from the crystal structure of pUb:UblR0RBR (PDB code 5N2W) were used following removal of the Ubl domain and adjoining linker coordinates (residues 1-83). Several linker sections absent in crystal structures of parkin were incorporated using the Modeller (Eswar *et al*, 2006) plug-in for UCSF Chimera (Pettersen *et al*, 2004). The tether (387-405) that partly occludes the RING1 binding site was allowed to move an average 5.3 Å in PyMOL (Delano, 2002). Coordinates for ubiquitin (PDB code 1UBQ) (Vijay-Kumar *et al*, 1987) and UbcH7 (PDB code 4Q5E) (Grishin *et al*, 2014) were used to create the UbcH7-Ub conjugate in the complex by using a single unambiguous restraint between the C-terminal G76 of ubiquitin and the catalytic C86K of UbcH7. An upper distance limit of 4.0 Å was set for ambiguous distance restraints while the unambiguous distance restraint was set to 6.8 Å. Standard parameters were used except inter_rigid (0.1) which was set to allow tight packing of the proteins, and the unambiguous force constants were set five-fold higher compared to those used the ambiguous constants. A total of 1,000 initial complexes were calculated and the best 100 structures were water-refined.

